# NanoNERF: A nanoscale NERF blaster replica made of DNA

**DOI:** 10.1101/2023.10.02.560388

**Authors:** Lauren P. Takiguchi, Mark B. Rober, Jocelyn G. Olvera, Amanda L. Wacker, Ryan J. Fantasia, Boyu Liu, Wade Shipley, Andrea Tao, Pallav Kosuri

## Abstract

We used DNA origami to create NanoNERF, the world’s smallest NERF blaster replica (Figure 1). We based our design on the NERF model *Maverick Rev-6*, and scaled the dimensions down three million times. NanoNERF is planar and measures ∼100 nm in length, with a length-to-width ratio closely resembling the original toy. Here, we describe the design, prototyping, and validation pipeline used to create the NanoNERF. We also discuss potential applications to motivate the creation of future nanoscale blasters with a firing functionality.

## INTRODUCTION

DNA origami creatively repurposes DNA as a structural building block, expanding the usefulness of this biological molecule beyond its conventional role of storing genetic information, and instead unlocking its potential for nanoscale construction^1-6^. DNA is a unique material due to its combination of biocompatibility, thermodynamic stability, and programmable assembly properties^3^. In particular, the ability to guide self-assembly through programmable base pairing enables predictable, base-pair resolution construction of intricate DNA nanostructures with unparalleled precision^4^.

DNA origami works by taking a long, single-stranded DNA scaffold with a known sequence and mixing it with hundreds of short, complementary staple strands that bind to and fold the scaffold into desired shapes (Figure 2, Supplementary Video 1). Thermodynamically driven to self-assemble through base-pairing interactions and stoichiometrically driven to optimize yield, these reactions can be very efficient with minimal user intervention^5^. A typical preparation of DNA origami in a drop of liquid results in trillions of fully folded, structurally similar objects.

**Figure 1:**
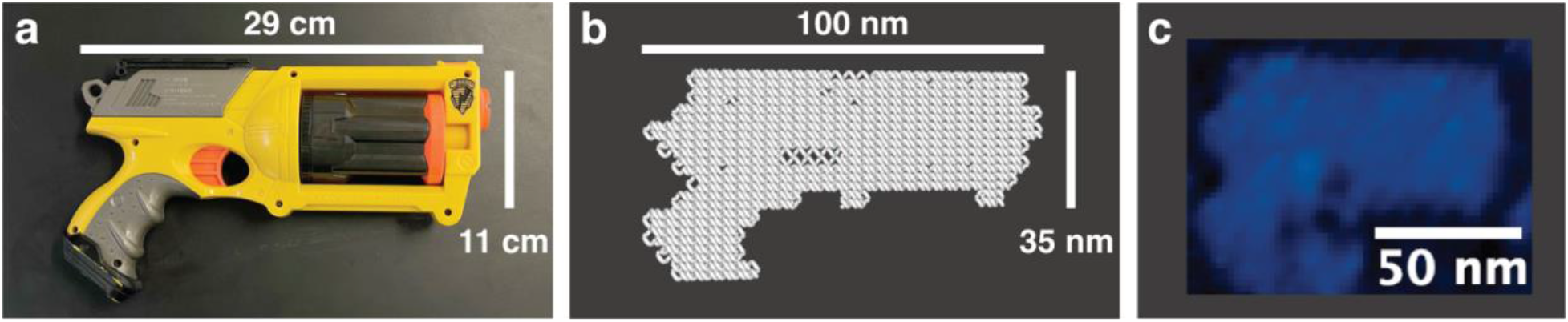
A nanoscale replica of the NERF Maverick Rev-6. a) Original NERF *Maverick Rev-6* toy, b) rendering of the NanoNERF structure in oxView. Gray lines represent individual DNA strands. NanoNERF length: 100 nm; width of barrel: 35 nm; thickness: 2 nm; c) Scan of a NanoNERF blaster acquired in an Atomic Force Microscope.

**Figure 2:**
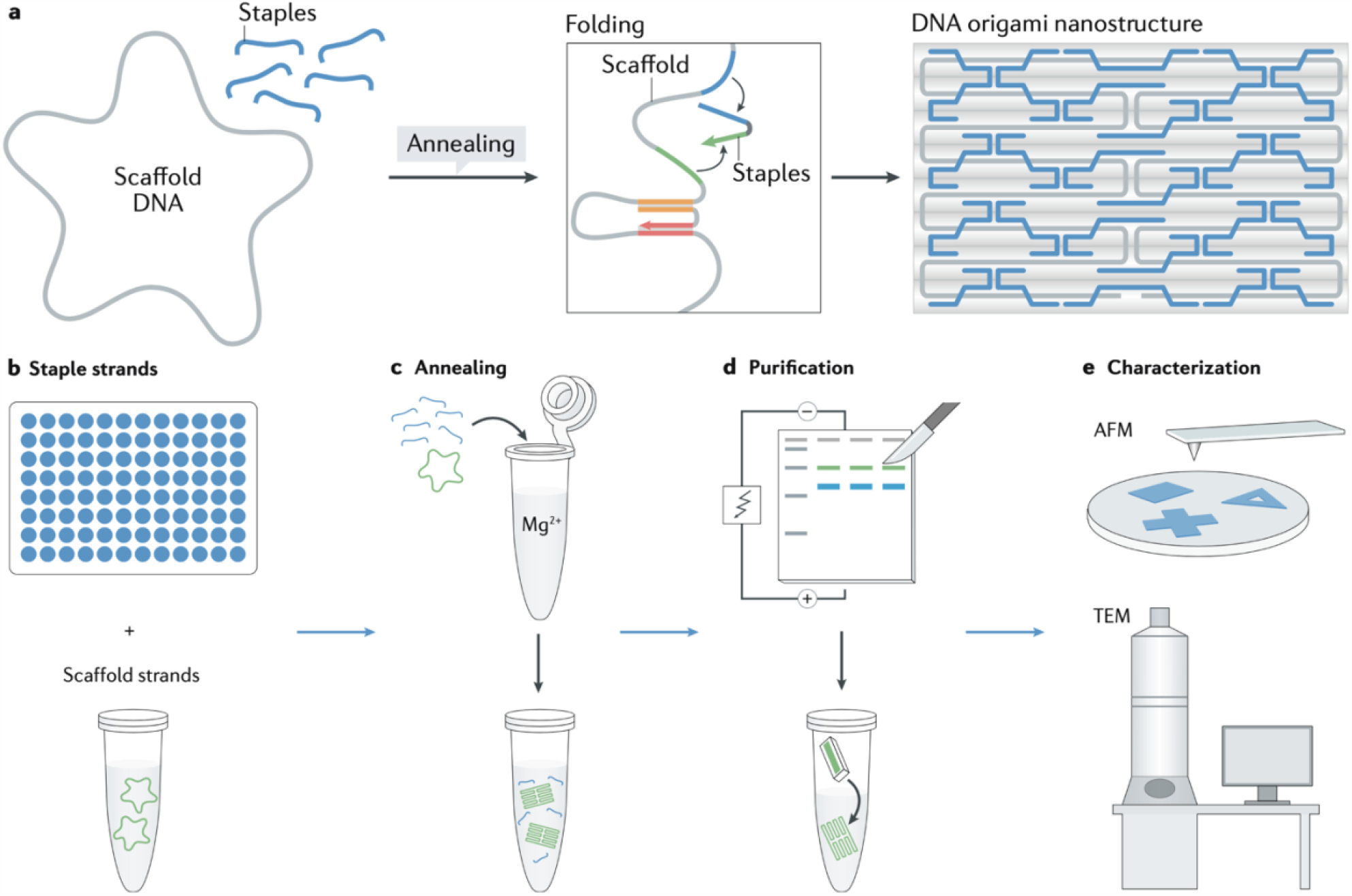
How DNA origami works. a) Process overview, b) Materials needed for self-assembly, c) Mixing scaffold strands and staple strands together for thermal annealing self-assembly reactions, d) Purification of DNA origami via agarose gel electrophoresis, e) Characterization of DNA origami with AFM and/or TEM. Adapted from ref. 3.

Whether constructing dynamic molecular machines, drug delivery vehicles, diagnostic devices, molecular breadboards, or nanoscale artwork, DNA origami offers engineers a creative canvas limited only by one’s imagination and the physical properties of DNA. Here, we demonstrate the nanoscale engineering capabilities of DNA origami by using it to create a miniaturized replica of a popular toy, the NERF blaster.

## RESULTS

The shape and proportions of our NanoNERF resemble the NERF *Maverick Rev-6* toy blaster. We designed the NanoNERF to have a length of 100 nm, a barrel width of 35 nm, and a thickness of 2 nm. We followed a standard DNA origami design protocol^6^ and used caDNAno^7^ for the strand routing and sequence design. Where possible, staples of length 35-45 bases were designed to have at least one 14-base “seed” region to nucleate assembly and maximize the yield of correctly folded objects. The routing of the scaffold and staples in our NanoNERF design were based on the new rectangular origami (NRO)^8^.

We used agarose gel electrophoresis and Atomic Force Microscopy (AFM) to validate successful folding of the NanoNERF (Figure 3). The gel scan in Figure 3a shows the scaffold alone (m13; left lane) and the NanoNERF samples folded in solutions with different concentrations of magnesium (lanes 2-5 from the left). Each of the NanoNERF lanes shows a single sharp band appearing above the level of the unfolded scaffold, indicating that in these origami samples, most of the scaffold strand had folded into a single species with a structure different from the scaffold alone – presumably the NanoNERF structure. Typically, folding of DNA origami is only successful within a narrow range of magnesium concentrations, however the NanoNERF bands appeared at the same vertical position in every lane, showing that the folding of the NanoNERF was not sensitive to magnesium concentration between 6 and 15 mM. The bright bands at the bottom of the gel contain the leftover staples that were present in excess as compared to the scaffold in each sample.

**Figure 3:**
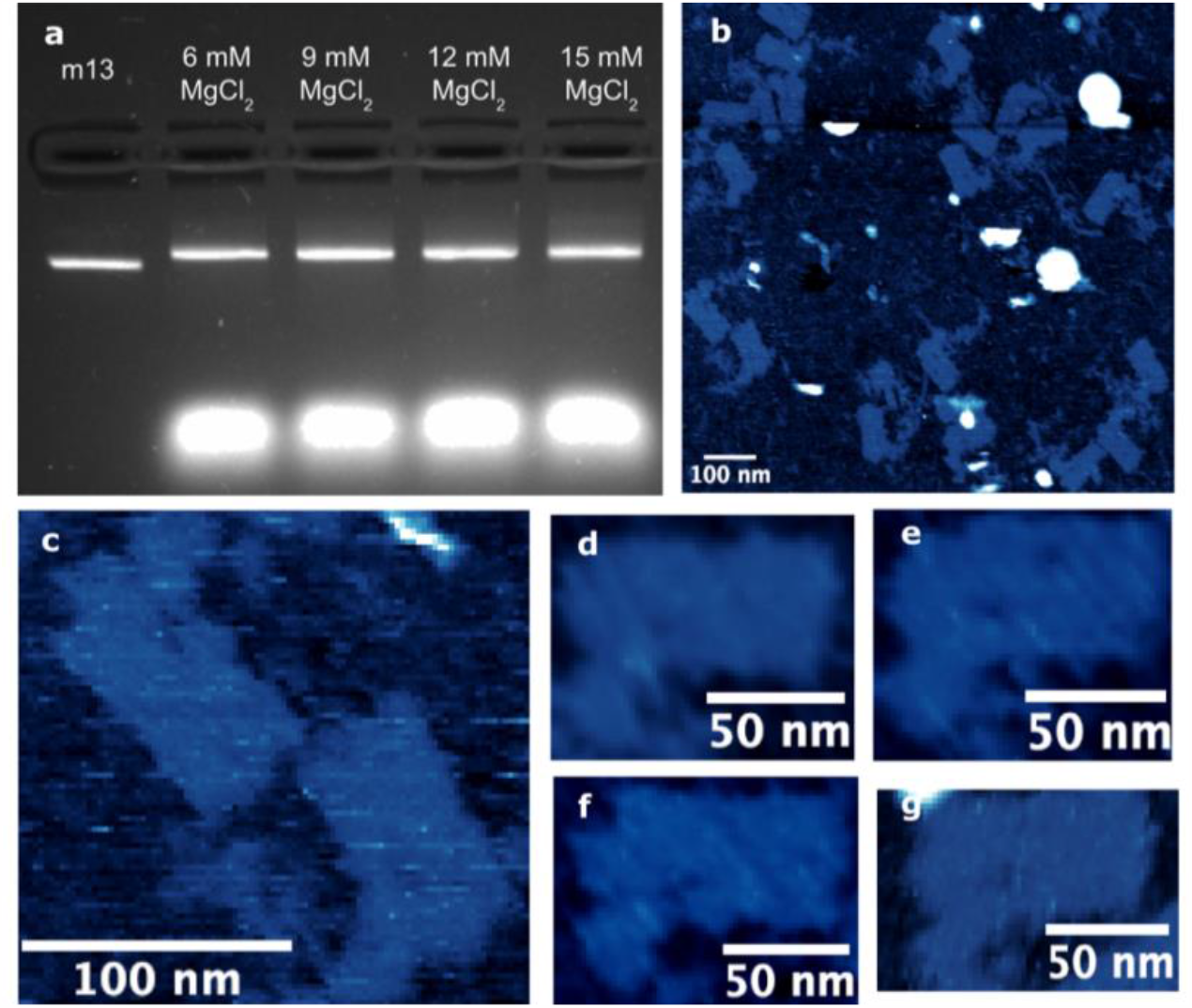
Results of NanoNERF folding. a) Image of agarose gel after electrophoresis of m13 scaffold (left lane) and NanoNERF samples folded in various magnesium concentrations (lanes 2-5), b-g) AFM scans of folded NanoNERF sample excised from gel.

After extraction of the folded sample from the gel, we imaged the NanoNERF objects in solution in an AFM. The AFM scans (Figure 3b-g) show clear NERF blaster shapes with the expected dimensions, showing that the folded species we identified in the gel corresponded to correctly folded NanoNERF blasters. We note that a few of the objects in the AFM scan do not appear to correspond to the correct structure. We speculate that these broken structures may be due to damage incurred during the AFM sample preparation or scanning process, rather than misfolding events, since the gel scan only showed a single sharp band containing the folded species – if the sample had contained a variety of different structures then the gel would likely have shown a smear rather than a sharp band.

## DISCUSSION

DNA origami is now more accessible to the public than ever before, both in financial cost and technical skill required^9^. With free, open-source software to aid each step of the design process, students and other aspiring engineers can begin the journey from behind their computer screen^7, 9-12^. Furthermore, the only laboratory equipment required to make a batch of DNA origami is a pipette and a programmable temperature bath, such as a standard thermocycler. We show an example DNA origami design pipeline in the methods section (Figure 4) that students and others can follow to bring their molecular designs to life.

**Figure 4:**
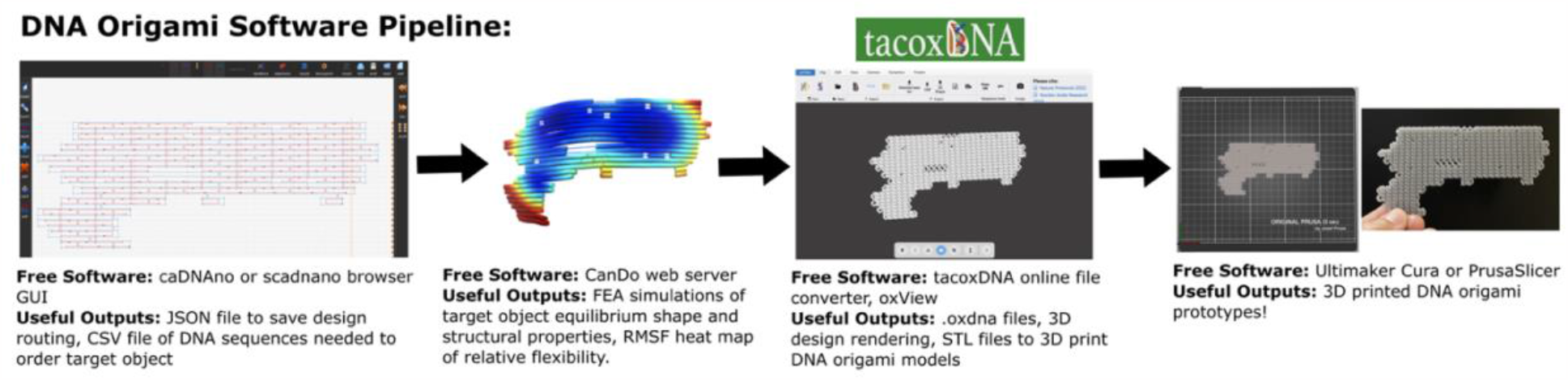
An open-source DNA origami design software pipeline. Flowchart showing some of the free software available to the public to start designing and creating DNA origami structures.

Here, we demonstrate the use of DNA origami to create replicas of NERF blasters at a scale of 1:3,000,000. To establish an intuition for the physical size of the NanoNERF, we asked the question: How many NanoNERFs could fit inside a spherical cell nucleus with a diameter of 10 μm? The volume enclosed by a 10 μm-sphere is (4 ⁄ 3) ∗ *πr*^3^ = 500 μm^3^ (500,000,000,000 nm^3^). By comparison, a NanoNERF occupies a volume of ∼10,000 nm^3^. Disregarding steric constraints, approximately 50 million (50,000,000) NanoNERFs could fit inside a single cell nucleus (see Supplementary Video 2 for a comparison of scales).

Unlike the original toy that fires foam darts, our NanoNERFs did not feature a firing mechanism. We propose that it would be possible to modify our design such that the NanoNERFs could fire a projectile, for instance by leveraging stored potential energy in the form of wound-up DNA^13^ or by using a different form of actuation. If we were to make functional NanoNERFs, what biological applications could we envision for such machines? One example of an application could be to inject DNA or RNA molecular cargo into cells, for instance to provide specific genetic instructions. DNA origami has recently been used for gene delivery to cells^14^. We speculate that the addition of a mechanical injection mechanism could improve the effectiveness of such devices. Supporting this point, we note that a protein-based molecular injection device was recently demonstrated for protein delivery to cells^15^. If coupled with a targeting mechanism (for instance by outfitting the NanoNERFs with antibodies), these blasters could be used to target cancer cells or pathogens for destruction.

Our NanoNERF provides an example of how we can translate engineering from the macroscopic to the nanoscopic world. There is surely no shortage of ideas for what can be done with DNA origami; there is just a shortage of scientists to execute them. We urgently need more scientists who are eager to tackle real-world challenges with new perspectives.

## MATERIALS AND METHODS

### DNA Origami Nanostructure Design

The NanoNERF was designed with caDNAno^7^ software (http://cadnano.org). The mechanical integrity of the structures were assessed using the CanDo^10^ software, a finite element analysis simulation that predicts equilibrium shape and physical properties of custom DNA nanostructures in solution (http://cando-dna-origami.org). The NanoNERF was designed with M13mp18 scaffold.

### Macroscopic 3D Printing of DNA Nanostructure Designs

Once a design was finalized in caDNAno, the file was exported as a .json file. TacoxDNA^11^ (http://tacoxdna.sissa.it/), a free online file converter, was used to convert the caDNAno .json file to oxDNA files. After the files were downloaded, the files were dropped in OxView^12^ (https://sulcgroup.github.io/oxdna-viewer/) and modified as necessary. In the “View” tab, the size of each of the “Visible Components’’ was increased ∼8 times, and the design was exported as a 3D shape (.gltf, .glb, .stl). Increasing the size of the components made the structure more robust to macroscopic 3D printing. The 3D shape file was imported into Ultimaker Cura slicing software, and a ∼10 cm prototype model was then printed on a Formlabs 3D printer.

### Self-Assembly Reactions

For scaffold, we used the m13mp18 single-stranded DNA (New England Biolabs). Staple strands were purchased from Integrated DNA Technologies (IDT), in 96-well plates and dissolved at 100 μM per oligo in a solution containing 10 mM Tris and 1mM EDTA buffered at pH 8 (see Supporting Information for oligo sets used in each nanostructure; sequences listed 5’ to 3’). The staple strands and scaffold strand were mixed in a folding buffer consisting of 10 mM Tris, pH 8.0, 1 mM EDTA, and 12 mM MgCl_2_. The concentrations of DNA in each reaction were 5 nM for the scaffold strand and 100 nM for the staple strands. Folding reaction mixtures were incubated and annealed using a thermocycler (Bio-Rad). To enable folding, the mixtures were held at 90 ºC for 15 minutes and annealed by cooling to 20 ºC in 1 ºC steps every 1 minute (for a total of ∼1.5 h folding time).

### DNA Origami Purification

Origami structures were purified from the folding mixtures using electrophoresis in a 2% agarose gel, run in an ice bath for 1 hour at 100 V, in running buffer consisting of 89 mM Tris pH 8.0, 89 mM borate, 2 mM EDTA, and 10 mM MgCl_2_. The appropriate bands were excised from the gel, and samples were extracted with Freeze N’ Squeeze spin columns (Bio-Rad) by centrifugation at 1,000 g for 60 minutes at 4 ºC. After centrifugation, the samples were concentrated using Amicon Ultra 0.5 mL, 100 kDa spin filters (Amicon) and spun for 12 min at 3,000 g at 4 °C. All samples were stored frozen at -20 °C until needed for further use. Note that the use of the agarose gel is important when folding a new origami structure to evaluate self-assembly products but is not necessary for later preps as samples can be purified with Amicon filters immediately after removal from the thermocycler.

### AFM Sample Preparation and Imaging

AFM images were obtained using a Bruker Innova AFM. A 5 μL droplet of purified origami sample and then a 30 μL drop of folding buffer with 10 mM NiCl_2_ added were applied to a freshly cleaved mica surface and left to incubate for at least 4 minutes. Images were acquired in liquid tapping mode, with the qp-BioT probe (Nanosensors).

## ACKNOWLEDGEMENTS

The authors acknowledge the use of facilities and instrumentation (AFM) supported by the National Science Foundation through the University of California San Diego Materials Research Science and Engineering Center DMR-2011924. We thank Amy Cao for creating the animations provided as part of the Supplementary Materials. We thank Steve Barry and Tony Artale for aiding in the 3D printing of the model NanoNERF. We are grateful for the coordination support provided by Roopal Rawani and Victoria Johnson, and for the creative vision of Mike Jeffs and Luke Hale.

## AUTHOR CONTRIBUTIONS

L.P.T., M.B.R., and P.K. conceptualized and supervised the project. L.P.T. designed the nanoNERF blaster and generated 3D models, with input from M.B.R. and P.K.. A.L.W., R.J.F., and B.L. folded and purified the NanoNERF. J.G.O. and W.S. acquired AFM images. L.P.T. processed AFM images. L.P.T. and P.K. wrote the manuscript with feedback from all authors.

## SUPPLEMENTARY MATERIALS

Online [Supplementary Materials](https://drive.google.com/drive/folders/18R0PDVmnPv47pBOVLAGyoiOmGmnaOwmo?usp=drive_link) include the NanoNERF caDNAno design (.json file), DNA sequences (listed 5’ to 3’), and oxView files used to make each structure. We also provide files used to create 3D-printed macroscopic prototypes of our design, and animations demonstrating DNA origami folding and the relative scales of the original NERF vs. NanoNERF.

